# Mode of inhibition of RNase P by gambogic acid and juglone

**DOI:** 10.1101/2022.10.20.512141

**Authors:** Nancy Wu Meyers, Agnes Karasik, Kipchumba Kaitany, Carol A. Fierke, Markos Koutmos

## Abstract

The first step in transfer RNA (tRNA) maturation is the cleavage of the 5’ end of precursor transfer RNA (pre-tRNA) catalyzed by ribonuclease P (RNase P). RNase P is either a ribonucleoprotein (RNP) complex with a catalytic RNA subunit or a <u>pro</u>tein-only <u>R</u>Nase <u>P</u> (PRORP). In most land plants, algae, and Euglenozoa, PRORP is a single-subunit enzyme. There are currently no inhibitors of protein-only RNase P that can be used as tools for studying the biological function of this enzyme. Therefore, we screened for compounds that inhibit the activity of a model PRORP from *A. thaliana* organelles (PRORP1) using a high throughput fluorescence polarization (FP) cleavage assay. Two compounds, gambogic acid and juglone (5-hydroxy-1,4-naphthalenedione) that inhibit PRORP1 in the 1 μM range were identified and analyzed. These compounds similarly inhibit human mtRNase P, a multi-subunit protein enzyme, and are 50-fold less potent against bacterial RNA-dependent RNase P. Biochemical measurements indicate that gambogic acid is a rapid-binding, uncompetitive inhibitor that targets the PRORP1-substrate complex while juglone acts as time-dependent inhibitor of PRORP1. X-ray crystal structures of PRORP1 in complex with juglone demonstrate the formation of a covalent complex with cysteine side chains on the surface of the protein. A model consistent with the kinetic data is that juglone binds to PRORP1 rapidly to form an inactive enzyme-inhibitor (EI) complex, and then undergoes a slow step to form an inactive covalent adduct with PRORP1. These inhibitors have the potential to be developed into tools to probe PRORP structure and function relationships.

## Introduction

Transfer RNAs are transcribed as precursor transcripts (pre-tRNA) with extra nucleotides at both the 5’ and 3’ ends and undergo several modifications before they are functional for use in protein synthesis. The first step in tRNA maturation across all domains of life is the removal of the 5’ leader through phosphodiester bond hydrolysis catalyzed by RNase P in the presence of Mg^2+^ (1–3). While RNase P is a ribonucleoprotein (RNP) consisting of a ribozyme and protein units in bacteria and in the nucleus of some eukaryotes (4–6), protein-only RNase P (PRORP) enzymes are present in the nuclei and organelles of eukaryotes (7–9). Nevertheless, both RNP and PRORP enzymes are proposed to employ a two metal ion mechanism for catalysis of the hydrolysis of a specific phosphodiester bond in pre-tRNA using a metal ion-bound water nucleophile (3, 10).

Human mitochondrial RNase P (composed of three protein subunits: MRPP1, MRPP2, and MRPP3) has been systematically shown to lack an RNA component (8). MRPP3 contains the active site for catalysis of pre-tRNA cleavage but requires the subcomplex composed of a R9 methyltransferase (MRPP1) and a short chain hydroxysteroid dehydrogenase (MRPP2) to activate function (11, 12). Disruption of the RNA processing pathway in mitochondria can be detrimental. Mutations of individual mitochondrial protein-only RNase P proteins in *D. melanogaster* are lethal, in part due to defective tRNA processing (13). siRNA knockdowns of either MRPP1 or MRPP3 in HeLa cells also result in mitochondrial pre-tRNA transcript accumulation (14). Furthermore, pathogenic mutations in MRPP1 and MRPP2 lead to dysfunctional mitochondrial RNA processing and decreased levels of respiratory chain complex proteins (11, 15). MRPP1 mutations were identified in two patients with birth defects. The fibroblast cells of these patients exhibit impaired mitochondrial RNA processing, increased levels of mitochondrial pre-tRNA transcripts and, as a result, defective mitochondrial protein synthesis of vital electron transport chain complexes (15). Furthermore, point mutations in mitochondrial tRNA (mt-tRNA) genes and mitochondrial RNase P proteins are associated with pathologies arising from mitochondrial dysfunction (16–19).

MRPP3, along with homologs including plant PRORP enzymes, belong to a unique class of endonucleases that contain a Nedd4-BP1, Yac Nucleases (NYN) domain, a member of the PIN domain-like superfamily (20, 21). PRORP1 and MRPP3 are similar in structure, with 22% sequence identity. In contrast to MRPP3, PRORP enzymes in most land plants, algae, and Euglenozoa do not require additional subunits for catalysis *in vitro* (7, 9, 22, 23). Land plants encode three functional paralogs of PRORP. Among these, the structure of *A. thaliana* PRORP1 and PRORP2 have been solved, revealing a pentatricopeptide repeat (PPR) domain that binds RNA and a structural zinc binding central domain in addition to the NYN domain (21, 24). Structures of N-terminal truncations of MRPP3 have also been solved, as well as a recent cryo-EM structure of the full MRPP1/2/3-pre-tRNA complex (25–27). Structural methods are valuable tools employed to understand the interactions within and between proteins and modulators. One study recently used X-ray crystallography to determine the binding site of purpurin on the protein subunit of *T. maritima* RNase P and identified juglone as an inhibitor (*IC*_50_ = 12.8 μM) of *T. maritima* RNase P, but the mode of juglone inhibition was unclear (28).

Researchers have traditionally monitored RNase P activity using radiolabeled RNA substrates coupled with denaturing urea-polyacrylamide gel electrophoresis (urea-PAGE) and autoradiography (29). To measure activity in real-time, fluorescence-based approaches have been introduced. A fluorescence polarization (FP) assay was employed using a fluorescently labeled pre-tRNA substrate to monitor activity and identify inhibitors of bacterial RNase P (30), and later used to monitor PRORP1 activity (3). In this assay, the substrate contains a 5’ fluorophore that is released upon cleavage by RNase P, which decreases the fluorescence polarization signal (3). Another approach using fluorescence resonance energy transfer (FRET) was introduced to identify small molecule inhibitors of bacterial RNase P (28). This method uses a minimal model pre-tRNA substrate with a 3’ fluorophore and a 5’ non-fluorescent quencher which are separated upon RNase P cleavage, resulting in an increase in fluorescence signal (28). Here we adapted the FP assay (3) to screen for inhibitors of PRORP1. PRORP1 is used as a model system to identify inhibitors and regulators of endonuclease activity and to better understand the mechanism of mitochondrial RNA processing *in vitro* and *in vivo*. There are currently no known inhibitors of protein-only RNase P enzymes. We screened for inhibitors using a small molecule compound library containing compounds extracted from microbes and pharmacologically active compounds. Dose-response assays validate that gambogic acid, a rapidly binding uncompetitive inhibitor, and juglone, a time-dependent inhibitor, are both at least 50-fold more potent against plant and human mitochondrial RNase P relative to *B. subtilis* RNase P. Additionally, juglone is about 10 times more potent against PRORP than *T. maritima* RNase P (*IC*_50_ = 12.8 μM) (28). Furthermore, we determined the site(s) of a covalent adduct between juglone and PRORP1 using X-ray crystallography. The compounds identified from this screen serve as lead compounds to probe mitochondrial RNase P activity *in vitro* and *in vivo*.

## Results

### A high throughput FP cleavage assay for multiple turnover (MTO) PRORP1 activity

We investigated the PRORP1 endonuclease activity using *B. subtilis* fluorescein-labeled pre-tRNA^Asp^ (Fl-pre-tRNA) that previously has been used to elucidate the catalytic mechanism of PRORP1 (3). To identify inhibitors of PRORP1, we adapted a FP cleavage assay that monitors the conversion of Fl-pre-tRNA to a fluorescein-labeled leader sequence cleaved by PRORP1 (Fig. 1). This assay has previously been used to measure cleavage catalyzed by PRORP1 and *B. subtilis* RNase P (3, 30).

**Figure 1.**
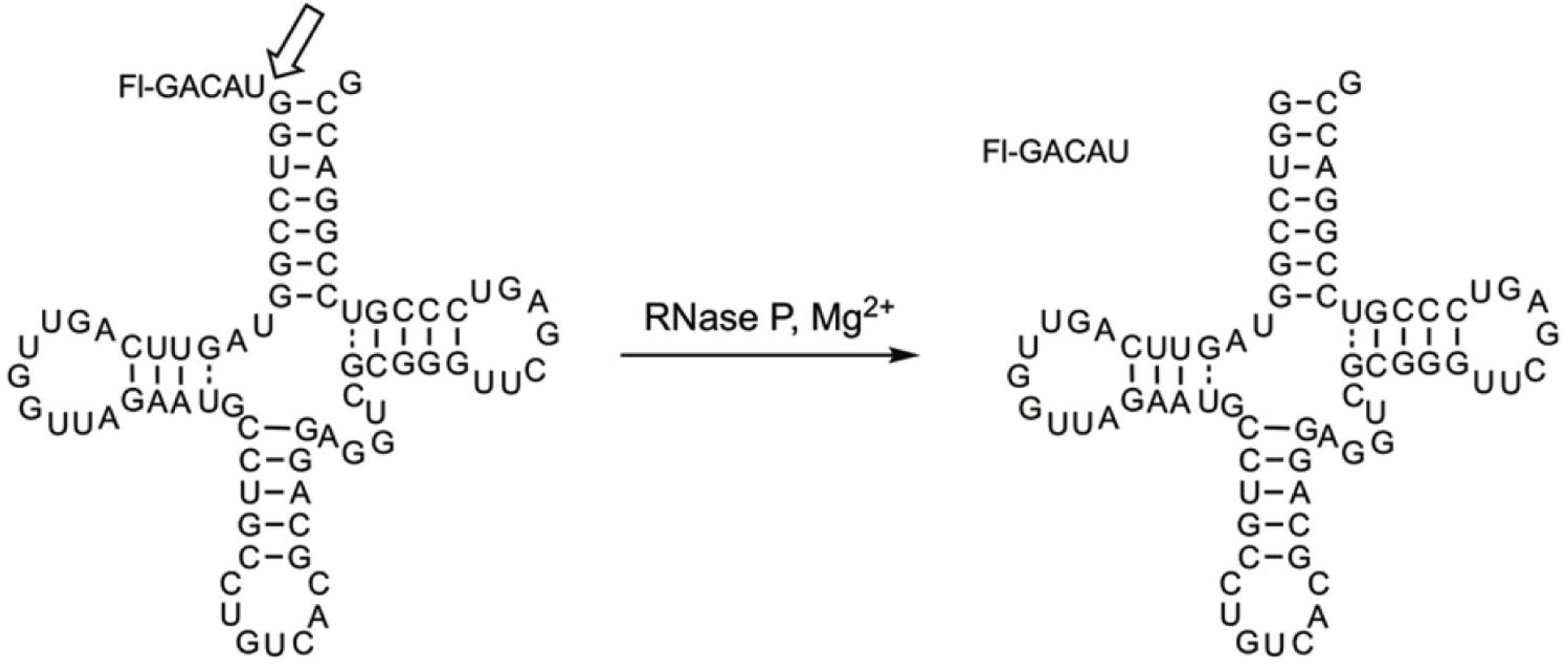
Fluorescence polarization (FP) assay to measure RNase P activity. Cleavage of fluorescein 5’ end-labeled *B.s.* pre-tRNA^Asp^ is catalyzed by RNase P to give products: fluorescein-labeled 5’ leader and tRNA^Asp^. The cleavage site is indicated by the open arrow.

We optimized this assay for use in a high throughput screen by varying buffer conditions to enhance PRORP1 activity, including the addition of spermidine and bovine serum albumin (BSA). We also added 500 nM yeast tRNA^mix^ to minimize false positive hits due to compounds binding to the pre-tRNA substrate. FP cleavage assays were performed using 800 nM Fl-pre-tRNA^Asp^, that corresponds to the *K*_M_ value therefore ensuring that both enzyme and enzyme-substrate species are present. This allows for the identification of both competitive and noncompetitive inhibitors, respectively. PRORP1 requires MgCl_2_ for catalytic activity. In the presence of CaCl_2_ PRORP1 does not catalyze cleavage, therefore CaCl_2_ was used to quench these reactions. A time course of PRORP1-catalyzed cleavage is shown in Fig. 2*A*. To determine the effectiveness of the FP cleavage assay we determined an average plate Z-factor of 0.6 (Fig. 2*B*), indicative of a robust assay (31). The relative inhibition activity of the positive and negative controls is shown in Fig. 2*C*.

**Figure 2.**
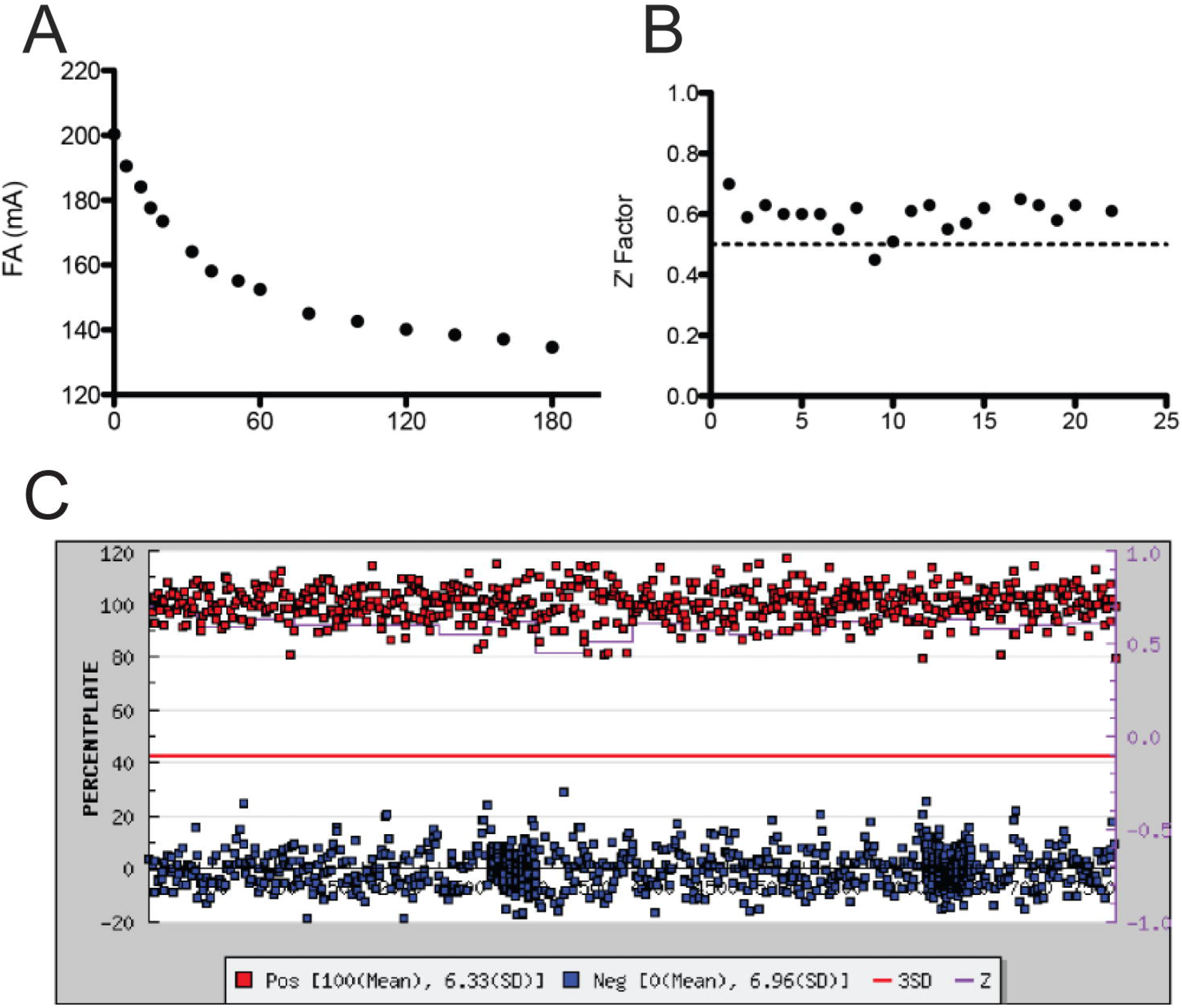
A robust high throughput FP cleavage assay for identifying inhibitors of PRORP1. *A*, a representative FP assay time course. Reactions were performed using 0.3 nM PRORP1 and 800 nM Fl-pre-tRNA^Asp^ at 22 °C, in 30 mM MOPS, pH 7.8, 1 mM TCEP, 5 mM MgCl_2_, 100 mM NaCl, 500 nM yeast tRNA, 5 mM spermidine, 6.65 μg/mL BSA, and 0.01% NP-40. Reactions were quenched by addition of 80 mM CaCl_2_ at various time points and the fluorescence anisotropy (FA) value was subsequently measured. *B*, the Z-factor values across each 384-well microplate range from 0.45 to 0.7. The average Z-factor of this assay is 0.6 (31). *C*, the percent inhibition of PRORP1 activity by a given compound in each plate was determined based on the FA value following 1 hr of reaction time. Values are shown for positive controls (addition of 80 mM CaCl_2_ before the reaction is started) in red and negative controls (1.25% DMSO with no inhibitor) in blue. The red line represents 3 standard deviations above the negative control.

Our optimized FP cleavage assay was used to screen a pilot compound library of small molecules (6046 compounds); this screen identified 88 compounds (using a concentration of 12.5 μM) that decreased activity more than 3 standard deviations (SD) compared to the negative control (1.5% hit rate). A more stringent second cut-off was introduced to identify compounds that reduce PRORP1 activity by more than 2.5-fold at 12.5 μM concentration resulting in the validation of 42 hits (see Table S1). Finally, a gel-based pre-tRNA cleavage assay was used to further eliminate false positive hits (Fig. S4). From the 42 hits, 23 of the identified compounds incubated at 12.5 μM with PRORP1 and pre-tRNA for 1 hr resulted in <10% product formation, as observed on urea-PAGE.

We then measured the inhibition of PRORP1 at four to six different inhibitor concentrations using our FP cleavage assay. The percent PRORP1 activity at 6.25 μM inhibitor is listed in Table S2. Of the compounds tested, gambogic acid and juglone (structures shown in Fig. 3), were selected for an in-depth kinetic analysis to determine their modes of inhibition based on: potency, whether they were known inhibitors of other enzymes, and commercial availability.

**Figure 3.**
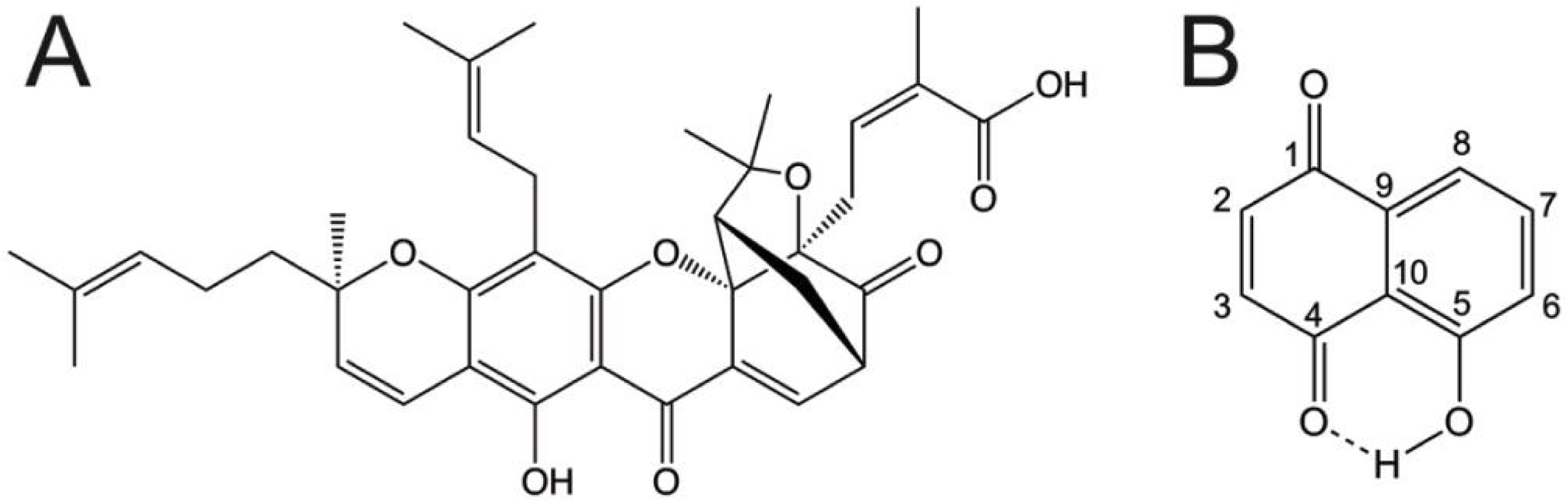
Structures of *A*, gambogic acid and *B*, juglone. *A,* the chemical structure of gambogic acid, a xanthonoid derived from the *Garcinia hanburyi* tree. *B,* the chemical structure of juglone, an allelopathic compound produced by plants of the *Juglandaceae* family.

### Gambogic acid and juglone are more potent inhibitors against PRORP1 and human mtRNase P than to RNP RNase P

To compare the potency of gambogic acid and juglone against PRORP and RNP RNase P, we determined the *IC*_50_ values of PRORP1, MRPP1/2/3, and *B. subtilis* RNase P using the FP assay. After a 30 min pre-incubation with PRORP1, gambogic acid inhibits PRORP1 activity with an *IC*_50_ value of 1.01 ± 0.04 μM and cooperative dependence (Hill coefficient, *n* = 4) and MRPP1/2/3 with an *IC*_50_ value of 0.26 ± 0.06 μM and no cooperative dependence (*n* = 1). Under these same conditions, juglone inhibits PRORP1 with an apparent *IC*_50_ value of 2.1 ± 0.04 μM (*n* = 1) and MRPP1/2/3 with an apparent *IC*_50_ value of 1.8 ± 0.04 μM (*n* = 1) (Fig. 4). Gambogic acid and juglone are both more than 50-fold less potent against *B. subtilis* RNP RNase P (*IC*_50_ value of gambogic acid is 50 ± 2 μM, *n* = 4; and *IC*_50_ value of juglone is greater than 500 μM, Fig. 4). These data suggest that gambogic acid and juglone recognize features of PRORP1 and MRPP1/2/3 that are not present in *B. subtilis* RNase P. This is unsurprising considering PRORP1 and MRPP1/2/3 are protein-only enzymes and do not have sequence homology with the P protein subunit of *B. subtilis* RNase P.

**Figure 4.**
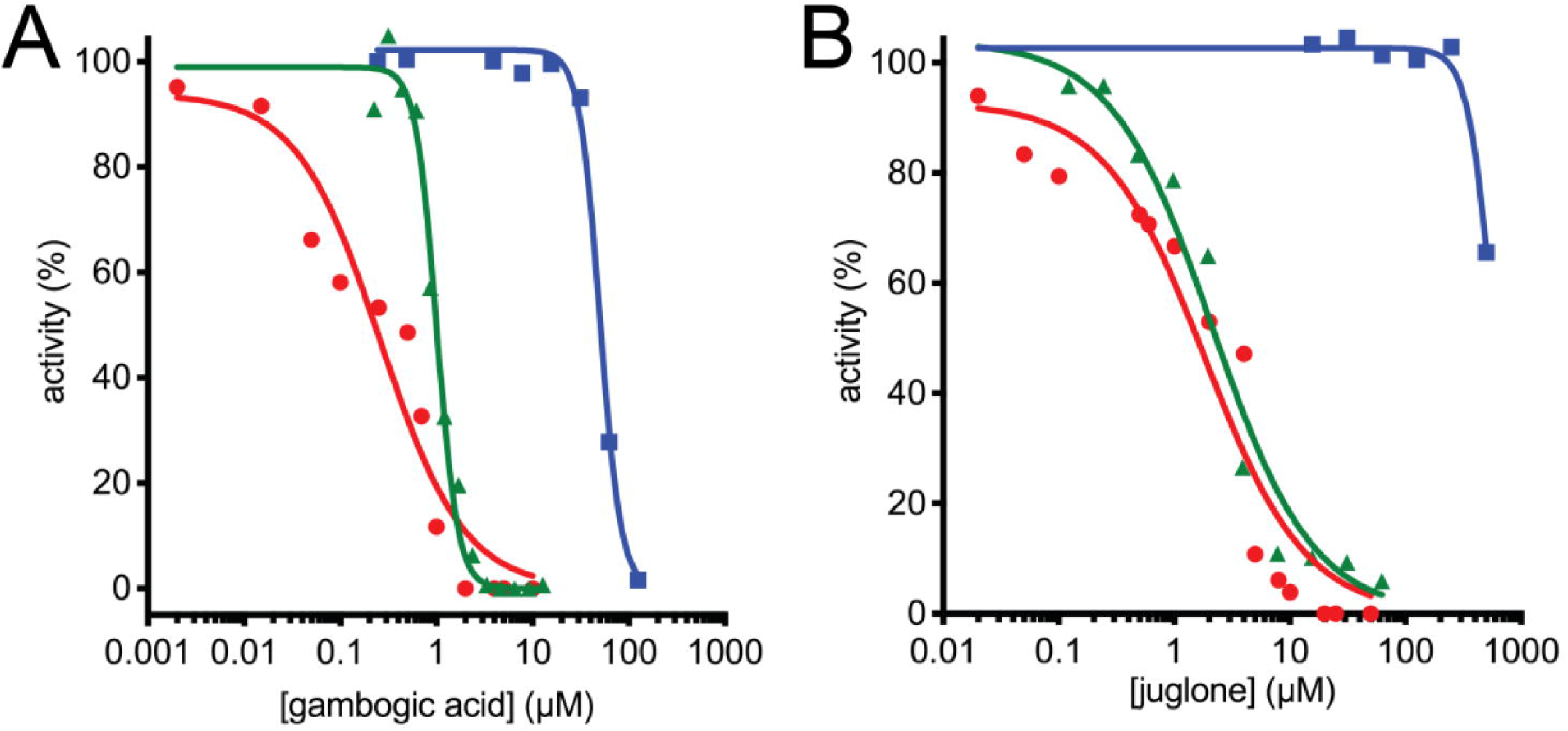
Gambogic acid (*A*) and juglone (*B*) inhibit *A. thaliana* PRORP1 (green), human MRPP1/2/3 (red), and *B. subtilis* RNase P (blue). Compounds were incubated with enzyme at double the reaction concentration for 30 minutes prior to the addition of an equal volume of substrate to initiate the reaction. PRORP1 reactions were performed with 0.3 nM PRORP1 and 800 nM pre-tRNA containing 40 nM Fl-pre-tRNA at 25 °C in 30 mM MOPS pH 7.8, 5 mM MgCl_2_, 100 mM NaCl, 1 mM TCEP, 500 nM (12 μg/mL) yeast tRNA, 5 mM spermidine, 6.65 μg/mL BSA, and 0.01% NP-40. MRPP1/2/3 reactions were performed using 50 nM MRPP3, 500 nM MRPP1/2, 500 nM pre-tRNA containing 40 nM Fl-pre-tRNA at 37 °C in 50 mM MOPS (pH 7.8), 1 mM TCEP, 4.5 mM MgCl_2_, and 100 mM NaCl. *B. subtilis* RNase P reactions were performed using 20 nM RNase P and 500 nM pre-tRNA containing 40 nM Fl-pre-tRNA at 37 °C in 50 mM Tris, pH 7.2, 5 mM MgCl_2_, 100 mM KCl, 500 nM (12 μg/mL) yeast tRNA, 20 mM dithiothreitol (DTT), 10 mM spermidine, and 0.01% NP-40. Equation 1 was fit to each dataset.

### Gambogic acid is an uncompetitive inhibitor of PRORP1

To gain a mechanistic understanding of gambogic acid and juglone, we investigated their modes of inhibition against PRORP1. There is no previous work on the detailed kinetic mechanism of these compounds. To obtain insights into the mode of inhibition of gambogic acid, we evaluated the time-dependence of PRORP1 inhibition using the FP assay. Pre-incubation of PRORP1 with gambogic acid prior to the addition of pre-tRNA to initiate the cleavage reaction does not affect the level of inhibition, consistent with a rapid equilibrium inhibition mechanism. Increasing gambogic acid concentration decreased the value of *k_cat_*_,app_ with no significant change in (*k*_cat_*/K*_M_)_app_ (Fig. 5), indicative of an uncompetitive inhibitor. A fit of equation 3 to these data provided a *k*_cat_ value of 1.86 ± 0.09 s^−1^ with an inhibition constant (*K*_I_) of 1.08 ± 0.05 μM and a Hill coefficient (*n*) of 4. This Hill coefficient for gambogic acid inhibition of PRORP1 suggests cooperative inhibition, multiple binding sites, or aggregation (32). A global fit analysis (Equations 2, 3, and 4) resulted in a similar *K*_I_ value (1.1 ± 0.03 μM). The data are consistent with an uncompetitive inhibition model where the inhibitor binds only to an ES complex (Scheme 1) and interacts with either or both the enzyme and bound pre-tRNA.

**Figure 5.**
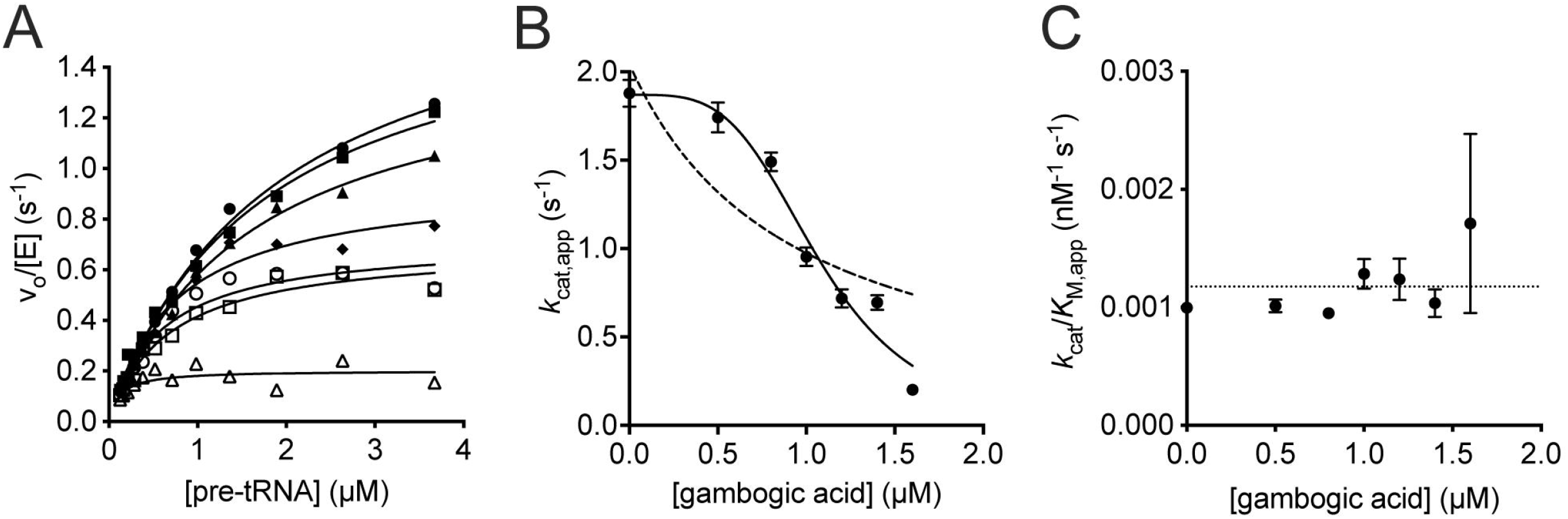
Gambogic acid is an uncompetitive inhibitor of PRORP1. *A*, the initial rate for PRORP1 catalyzed cleavage of Fl-pre-tRNA is plotted as a function of substrate concentration over a range of concentrations of gambogic acid (filled circles, 0 μM; filled squares, 0.5 μM; filled triangles, 0.8 μM; filled diamonds, 1 μM; open circles, 1.2 μM; open squares, 1.4 μM; open triangles, 1.6 μM). Equation 2 was fit to each data set to calculate the values of *k*_cat,app_, and *K*_M,app_, and (*k*_cat_/*K*_M_)_app_. *B, k*_cat,app_ values for PRORP1 activity are obtained from fitting Equation 2 to the data in A. Equation 3 is fit to the dependence of *k*_cat,app_ on gambogic acid concentration, resulting in an inhibition constant (*K*_I_) of 1.1 ± 0.05 μM and a hill coefficient (*n*) of 3.8 ± 0.8. The dotted line simulates the fit of equation 3 with *n* = 1. *C, k*_cat_/*K*_M,app_ values for PRORP1 activity vary little as a function of the gambogic acid concentration. Error bars indicate standard errors obtained from fitting the data.

**Scheme 1.**
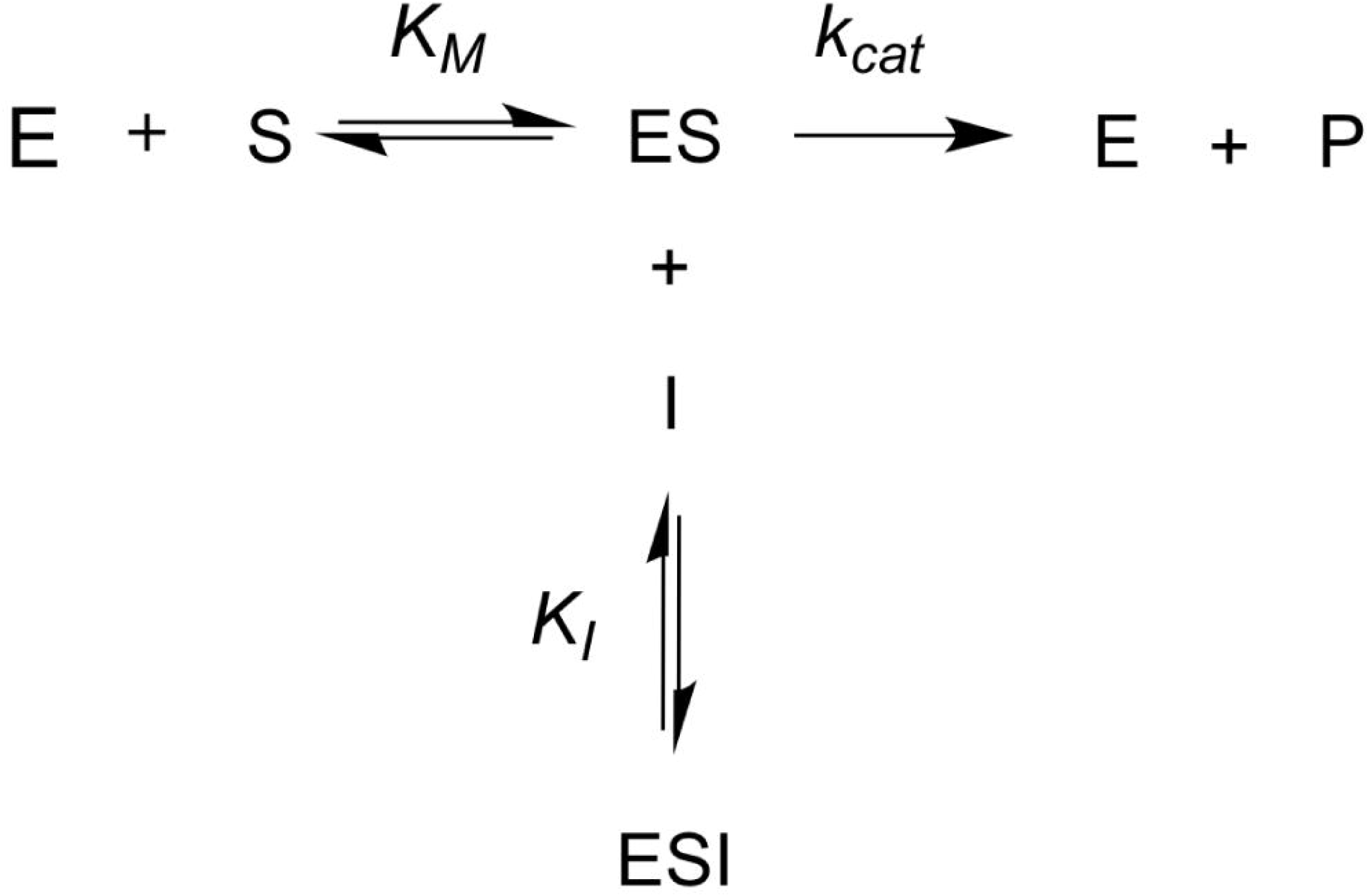
An uncompetitive inhibitor (I) binds exclusively to the enzyme-substrate (ES) complex to inhibit the enzyme. The inhibitor decreases the value of *k*_cat_ but does not affect *k*_cat_/*K*_M_ values since it does not bind to the enzyme without the substrate. For gambogic acid, the cooperativity of the inhibition data suggests that multiple molecules of gambogic acid are involved in the mechanism of inhibition.

### Juglone is a time-dependent inhibitor of PRORP1

Inhibition of PRORP1 by juglone was dependent on the length of the pre-incubation time. PRORP1 activity decreased with increasing pre-incubation time following a single exponential decay (Fig. 6*A*). Extrapolation of the activity to 0 min pre-incubation revealed an initial rapid inhibition step (Fig. 6*B*) prior to a slow inactivation step. The initial activity decreases as the concentration of juglone increases with a 50% decrease in activity at the highest concentration (10 μM) of juglone. Furthermore, the rate constant for inactivation (*k*_inact_) of PRORP1 is linearly dependent on the concentration of juglone (Fig. 6C). These data were used to estimate the apparent inhibition constant (*K*_I_) and the value of *k*_inact_*/K*_I_. A fit of Equation 1 to the dose response curve of the activity extrapolated to 0 min pre-incubation data provides an estimate of the apparent initial inhibition constant (*K*_I_ = 12.5 ± 2.7 μM) (Fig. 6*B*); here the *IC*_50_ value approximates the *K*_I_ value since the substrate concentration (0.25 μM) is below the *K*_M_ value (0.8 μM, Fig. S1). The solubility of juglone limited our ability to measure the time-dependent inhibition at concentrations above the estimated *K*_I_ value. The observed pseudo first order rate constant (*k*_inact,app_) was estimated by fitting Equation 5 to each of the datasets in Fig. 6*A*. A hyperbolic fit of the data using equation 6 estimates the *k*_inact_*/K*_I_ value of 0.0010 ± 0.0001 μM^−1^ min^−1^ (Fig. 6*C*). Additionally, the value of *k*_inact_*/K*_I_ (0.001 ± 0.00004 μM^−1^ min^−1^) was calculated from a global fit of all the data in Fig. 6*A* using Equations 5 and 6. These data are consistent with a two-step inhibition mechanism where juglone rapidly binds to and inhibits PRORP1, followed by a slow conversion from the enzyme-inhibitor complex (EI) to a second EI* complex described by *k*_inact_ (Scheme 2).

**Figure 6.**
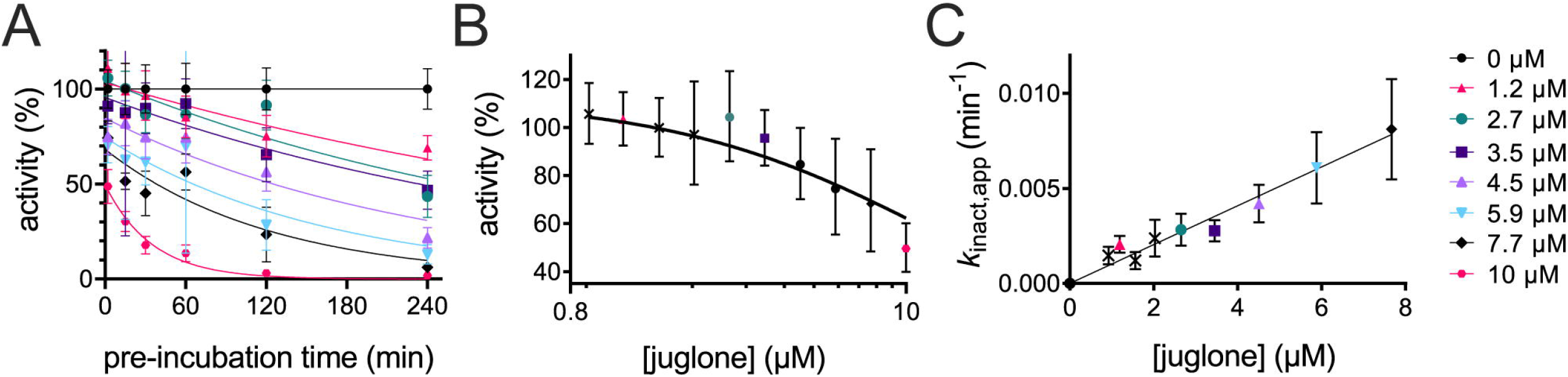
Juglone is a time-dependent inhibitor of PRORP1 exhibiting a 2-step inhibition mechanism. *A*, the initial rate of PRORP1-catalyzed cleavage decreases with increasing pre-incubation with juglone (0 – 10 μM). A global fit of the data (Equations 5 and 6) was used to calculate *k*_inact_ /*K*_I_. *B*, each dataset in A was extrapolated to 0-min pre-incubation and Equation 1 was fit to the data to calculate *IC*_50_, which is equal to the *K*_I_ (12.5 ± 2.7 μM, *n* = 1). *C*, the observed pseudo first order rate constant (*k* _inact,app_) for the inactivation of PRORP1 was fit using Equation 6 and exhibits a linear dependence on juglone concentration below the *K*_I_ value. The errors shown are the standard errors obtained from fitting Equation 5 to the data in *A* independently. Reactions were performed with 0.3 nM PRORP1, 250 nM pre-tRNA containing 40 nM Fl-pre-tRNA, at 22 °C, in 30 mM MOPS, pH 7.8, 1 mM TCEP, 5 mM MgCl_2_, 500 nM (12 μg/mL) yeast tRNA, 5 mM spermidine, 6.65 μg/mL BSA, and 0.01% NP-40. Data is fit to determine kinact/KI. It does not look hyperbolic because we are well below Ki.

**Scheme 2.**
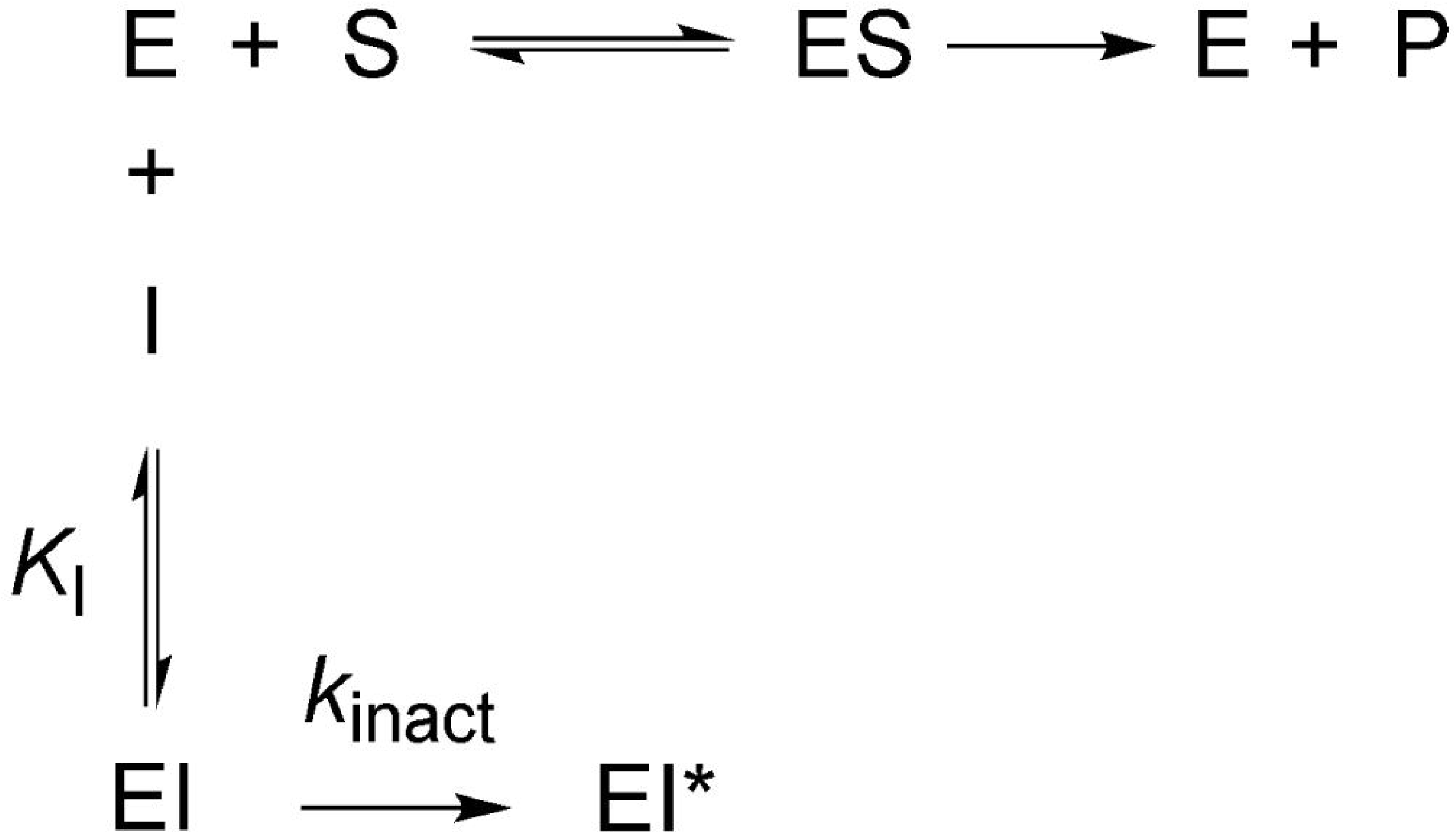
Model of 2-step inactivation mechanism of a slow-binding inhibitor. Juglone (represented by I) inactivates PRORP1 (represented by E) through a rapid first step to form EI described by *K*_I_, followed by a slow second step to form EI* described by *k*_inact_. EI and EI* are both inactive species of the enzyme.

### Juglone covalently modifies surface-exposed cysteine residues in PRORP1

To understand the molecular basis for the time-dependent inhibition of PRORP1 by juglone, PRORP1 was crystallized as previously described (19), in the absence of inhibitor and subsequently the crystals were soaked for varying times in juglone-containing solutions. Since crystallization conditions are different from our assay conditions, we optimized the time course of juglone-PRORP1 soaking experiments. PRORP1 crystals were incubated with juglone for 45 minutes, 3 hours and overnight. Upon soaking the transparent PRORP1 crystals in juglone, the crystals turned dark yellow, indicative of juglone incorporation. The resulting PRORP1-juglone soaked crystals diffracted between 1.8-2.2 Å (. In all solved structures, PRORP1 was covalently modified by juglone at C353, which is located in a solvent accessible, flexible region between the metallonuclease and zinc-binding central domains making several stacking and polar interactions with surrounding side chains (Fig. 7). Observed electron densities for the C353-juglone conjugate indicate full occupancy for this site in all crystal structures (Fig. 7*B*) and juglone B factors are comparable with B factors of surrounding residues. While crystals exposed to juglone for 45 minutes contain only C353 modified by juglone (PDB ID: 6BV5), structures based on crystals soaked for 3 hours (PDB ID: 6BV6) contained two additional modification sites (C373 and C346) albeit these sites were not fully occupied (Fig. S2). In addition to the presence of modification in C373, overnight soaking with juglone revealed an additional modification in the PPR domain, at residue 222 (PDB ID: 6BV9). However, only partial juglone electron densities were observed at C373 and C222 for the overnight soaked crystals (Fig. S2). Although multiple juglone modification sites were observed, structural rearrangements in the protein because of these modifications were not evident.

**Figure 7.**
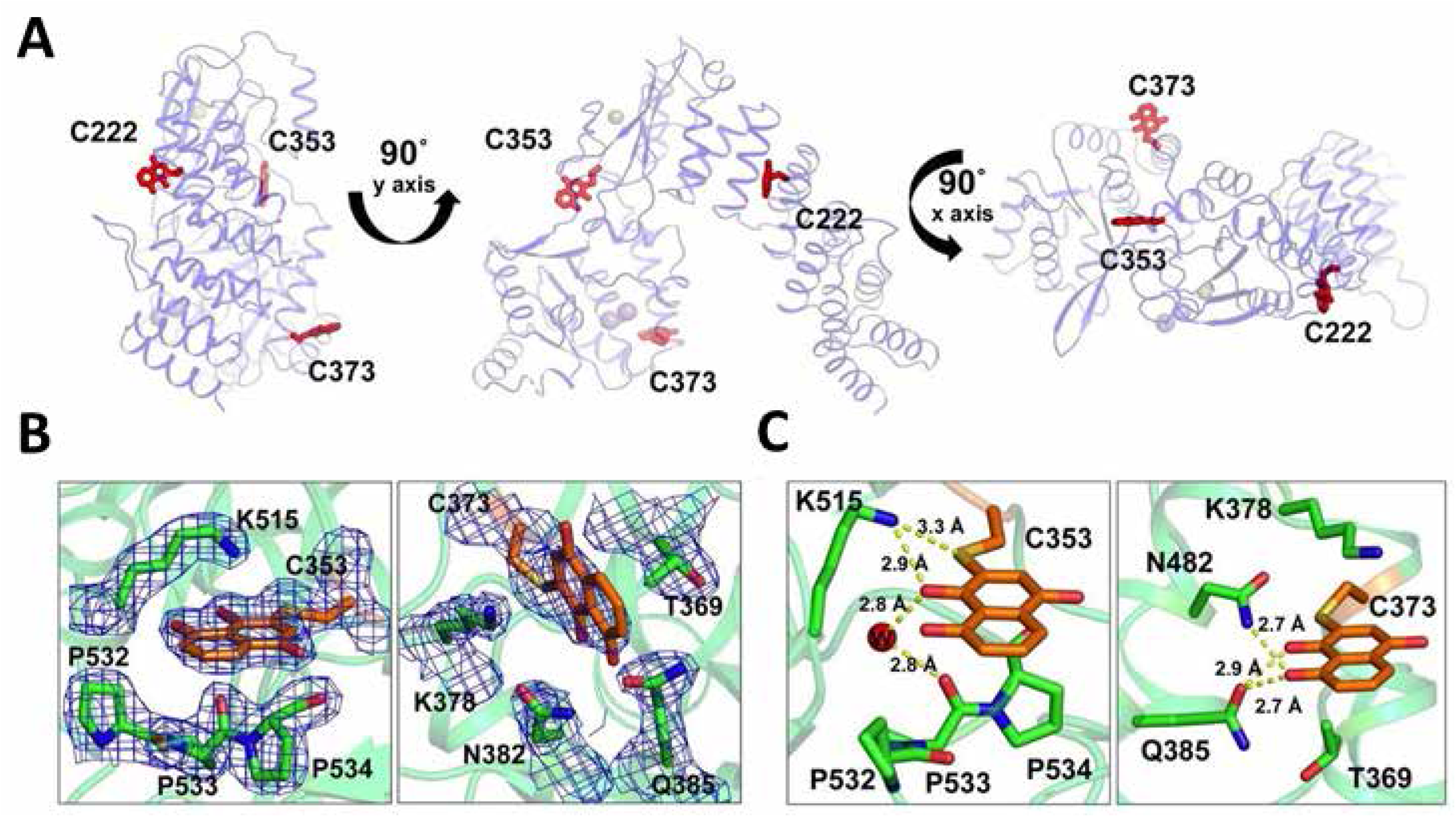
Crystal structure of PRORP1 in conjugation with juglone after overnight incubation. *A*, PRORP1 makes a covalent interaction with juglone (atoms shown in sticks and C colored orange), through cysteine thiol side chains at C222, C353, and C373. Active site metals were modeled from a previously solved structure of WT PRORP1 (4G26) with Mn^2+^ ions shown as spheres. *B*, electron densities and *C*, interactions of juglone-cysteine conjugates (C 353 and 373) are shown with surrounding residues.

We also addressed whether the mode of inhibition of juglone by selective cysteine modification in PRORP1 would change in the presence of catalytic metal ions. Hence, we soaked PRORP1 crystals with Mn^2+^ metal bound to the active site with juglone. The bound Mn^2+^ did not affect juglone modification of the previously identified sites and vice versa, the presence of juglone did not alter metal binding at the active site (PDB ID: 6BV5 and 6BV8).

Given the multiple sites of cysteine modification we wanted to identify the one(s) that potentially play a significant role in PRORP1 inactivation. Therefore, several cysteine-to-alanine point mutants were expressed, purified and subjected to inhibition studies. We tested both single-site (C353A and C373A) and double-site (C353A/C373A and C222A/C353A) mutations. Time-dependent inactivation by juglone was observed for each of the mutants tested (Fig. 8*A*). The value of *k*_inact,app_ for each PRORP1 at varied concentrations of juglone was determined by fitting a single exponential equation (Equation 5) to the data (Fig. 8*A*). Unexpectedly, the values of *k*_inact_ /*K*_I_ (reflecting the inhibition of E + I to form EI*) increased for a number of the PRORP1 variants: wild-type (0.001 ± 0.00004 μM^−1^ min^−1^), C353A (0.005 ± 0.002 μM^−1^ min^−1^), C373A (0.002 ± 0.0004 μM^−1^ min^−1^), C222A/C353A (0.003 ± 0.0001 μM^−1^ min^−1^), and C353A/C373A (0.0008 ± 0.0002 μM^−1^ min^−1^). These values were determined from fitting Equation 6 to each of the datasets shown in Fig. 8*B*. These modest increases could be due to a decrease in the *K*_I_ value or an increase in the value of *k*_inact_. Over the range of concentrations measured, little curvature was observed in the dependence of *k*_inact_ on the juglone concentration and modest to no initial inhibition was observed (Fig. 8*B*, Fig. S3), suggesting that the slight increase in *k*_inact_ /*K*_I_ is due mainly to an increase in *k*_inact_. Altogether, these data demonstrate that juglone reacts with multiple exposed cysteine sites with C353 as the kinetically preferred site in the crystals. Although, we cannot discount the possibility, however unlikely, that juglone may employ a mode of inhibition beyond cysteine residue modification that our crystal structures fail to capture.

**Figure 8.**
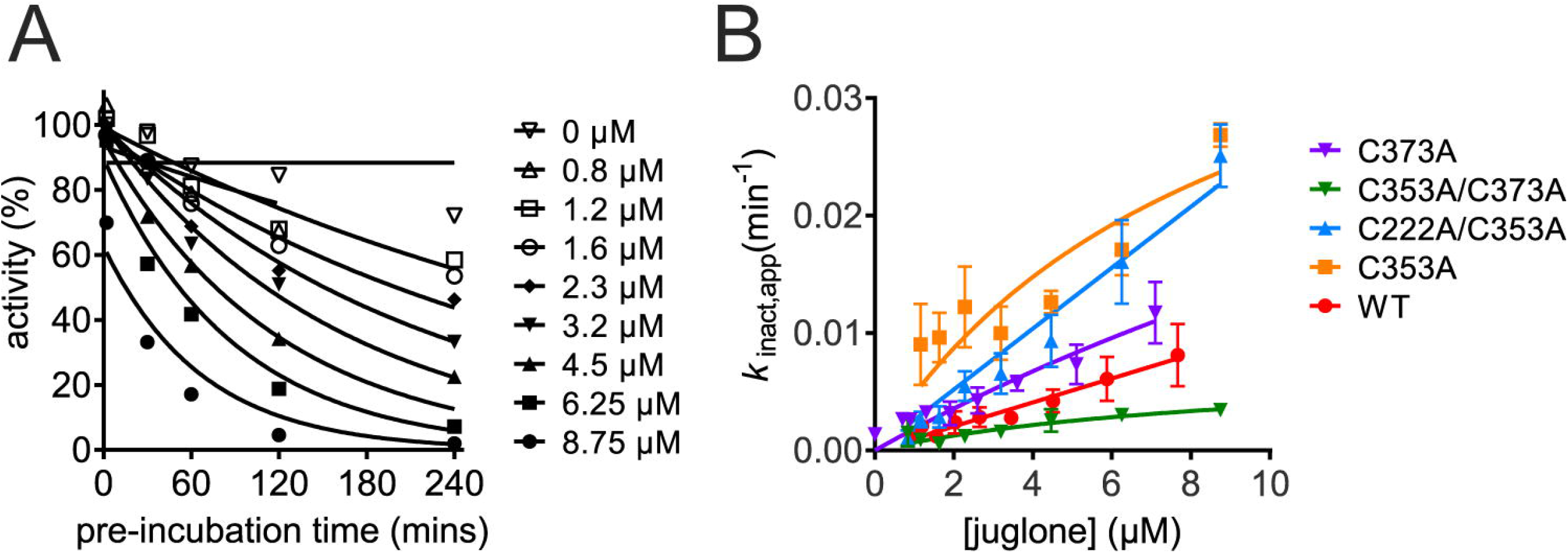
Effects of PRORP1 cysteine mutations on juglone inactivation. *A*, the PRORP1 C353A activity at varying juglone concentrations is plotted against the pre-incubation time. A fit of Equation 5 was used to determine the value for *k*_inact_. *B*, values of *k*_inact_/*K*_I_ were determined for each PRORP1 variant: wild-type (red, 0.001 ± 0.00004 μM^−1^ min^−1^), C353A (orange, 0.005 ± 0.002 μM^−1^ min^−1^), C373A (purple, 0.002 ± 0.0004 μM^−1^ min^−1^), C222A/C353A (blue, 0.003 ± 0.0001 μM^−1^ min^−1^), and C353A/C373A (green, 0.0008 ± 0.0002 μM^−1^ min^−1^) by fitting Equation 6 to the data. Error bars indicate standard errors obtained from fitting the data. Reactions were performed using the conditions listed in the caption of Fig. 6.

## Discussion

We have identified two potent inhibitors against PRORP1 and MRPP1/2/3 and conducted an extensive investigation of their modes of inhibition against PRORP1. Additionally, we report the first structural study relating to juglone’s potential universal mode of inhibition. Our FP assay identified compounds from a small library (6046 compounds) with different inhibitory mechanisms. Kinetic analysis revealed that gambogic acid inhibits PRORP1 with bound pre-tRNA; this inhibitor (shown as I) binds to the PRORP1/pre-tRNA complex reversibly, leading to a decrease in both *K*_M,app_ and *k*_cat,app_ (Fig. 5). The data are consistent with the behavior of a reversible uncompetitive inhibitor (Scheme 1). Gambogic acid inhibits PRORP1 activity with cooperativity (*n* = 4), which suggests multiple binding sites. Gambogic acid is an uncompetitive inhibitor that binds to the PRORP1/pre-tRNA complex so it could interact with multiple sites on PRORP1, the PRORP1/pre-tRNA interface and/or bound pre-tRNA. The lack of cooperativity (*n* = 1) in inhibition of MRPP1/2/3 is inconsistent with gambogic acid aggregation. The difference in cooperativity suggests differential binding to PRORP1 compared to MRPP1/2/3, despite the homology between the two proteins. Structural and enzymatic studies have indicated that binding of PRORP to pre-tRNA is followed by a conformational change that leads to a catalytically active structure (33, 34). One mechanism of inhibition consistent with the uncompetitive nature of gambogic acid is inhibition of this conformational change through ligand binding to the initial enzyme-substrate complex. The additional subunits in MRPP1/2/3 may stabilize alternate conformations of the bound pre-tRNA complex leading to the observed change in cooperativity. Gambogic acid binds to and stabilizes the PRORP1-pre-tRNA complex and this inhibitor may aid in obtaining the first co-crystal structure of PRORP1 with pre-tRNA.

Gambogic acid has many cellular targets ranging from kinases to proteins involved in maintaining the structural integrity of the cell and is thus considered a broad-spectrum clinical agent (35). It also has anti-inflammatory, anti-cancer, antibacterial, and antioxidant effects due to its ability to modulate many cell signaling pathway intermediates including epigenetic regulators, protein kinases, transcription factors, inflammatory biomarkers, and growth regulators (36, 37). Gambogic acid has been shown to inhibit a number of targets, but cooperative inhibition has not been observed with these targets. One study measuring the *IC*_50_ of gambogic acid and its analogs against gap junction intercellular communication (GJIC) did not observe cooperativity (38), similar to the result for MRPP1/2/3.

We also investigated the mode of inhibition of juglone and found that it reacts with surface-exposed cysteine residues on PRORP1, potentially affecting the enzyme’s ability to adopt a full range of structural conformations in the presence of pre-tRNA. Our structural work indicates that juglone can form a covalent bond with at least four surface-exposed cysteine residues in PRORP1 at the C-3 position of juglone, likely through a Michael addition reaction, as previously reported (39). Quinones such as juglone are potent electrophiles that readily react with thiol groups in proteins. We found that juglone exhibits the features of a slow binding inhibitor of PRORP1. Juglone binds to PRORP1 to form an inactive enzyme-inhibitor (EI) complex, which then undergoes a slow step to form another inactive EI* complex via formation of a covalent adduct with PRORP1 and/or a conformational change in PRORP1. Considering that no structural rearrangements were observed in the PRORP1 structure, we propose that the formation of the EI* complex reflects formation of the covalent complex. To understand the mechanism of juglone inactivation of PRORP1 and the role of reaction with cysteine residues, single and double point mutations were made and the inhibition activity was evaluated. Single-site cysteine mutations did not abolish inhibition activity, rather they generally increased the value of *k*_inact_ /*K*_I_ (Fig. 8). The C353A/C373A double mutation modestly reduced the value of *k*_inact_ /*K*_I_ compared to wild-type PRORP1. These mutagenesis data indicate that inactivation by juglone is not specific to reaction with the cysteine side chains on PRORP1 that were juglone conjugated in our structure.. Multiple sequence alignment of PRORP1 homologs identifies three conserved cysteines that are solvent exposed (Fig. S5). These residues (C344, C347 and C565) coordinate a Zn^2+^ ion within the zinc-binding site of the central domain, which plays a key role in structural stability and domain orientation. The essential role of these residues makes them potential specific sites of inhibition through juglone conjugation and is exemplified by these residues’ intolerance to mutation. Overall, our findings reveal two potential mechanisms of juglone inhibition. One where juglone non-specifically modifies surface exposed cysteines and above a certain threshold these modifications become deleterious to protein stability and function. The other, where juglone conjugation of specific structural cysteines impairs its function.

We used a FP high throughput assay to identify inhibitors that target PRORP1 and human mtRNase P with higher potency than bacterial RNase P. The potency of gambogic acid and juglone against PRORP1 is comparable with that against human mtRNase. However, the contribution of the MRPP1/2 subcomplex to mtRNase P activity could alter the mode of inhibition of these and other inhibitors, possibly by interacting with pre-tRNA bound to MRPP3. The ability of both gambogic acid and juglone to inhibit human mtRNase P demonstrates the promise of this approach for the identification of other potent inhibitors from a larger chemical library. Potent human mtRNase P inhibitors will be useful tools needed to identify the multiple functions of this enzyme in the cell. Inhibitors generally abolish enzyme activity more quickly and completely compared to siRNA knockdowns allowing for rapid identification of human mtRNase P substrates (14). Using the methods developed in this study, future work would include using human mtRNase P to screen a larger library of compounds to identify inhibitors for *in vivo* studies.

## Conclusion

We have demonstrated the use of a FP cleavage assay in a high throughput screen to identify compounds that inhibit PRORP1 and human mitochondrial RNase P activity. These compounds (gambogic acid and juglone) target different steps in PRORP1 activity. Gambogic acid is an uncompetitive inhibitor targeting the PRORP1-pre-tRNA complex while juglone is a time-dependent inactivator that forms a covalent complex with cysteine side chains. This work presents the first crystal structure of a protein that is covalently modified by juglone and identifies surface cysteine residues of PRORP1 that react with juglone. As we are developing tools to study human mtRNase P activity *in vitro*, future work would include translating this work to identify inhibitors with selectivity *in vivo*.

## Experimental Procedures

### Reagents

Buffer components were all purchased through Sigma unless specified. Baker’s yeast tRNA^mix^ was purchased from Roche. Gambogic acid was purchased from EMD Millipore and juglone (5-hydroxy-1,4-naphthoquinone) was purchased from Sigma at the highest purity. Recombinant His_6_-T7 RNA polymerase was expressed in *E. coli* and purified as previously described (40). Guanosine 5’-monothiophosphate (GMPS) was synthesized from 2’,3’-isopropylidene guanosine and thiophosphoryl chloride as previously described (41).

### Protein expression and purification

*A. thaliana* PRORP1 was expressed and purified according to literature procedures (21). The protein component of RNase P from *B. subtilis* was prepared as previously described (42). The MRPP3 gene (*KIAA0391*) with an N-terminal truncation (residues 46-583, Δ45MRPP3) was cloned into a pETM-11 vector that adds an N-terminal His_6_-tag as previously described (43). The plasmid was transformed and expressed in Rosetta (DE3) *E. coli* cells. Cells were grown in LB medium with 50 mg/L kanamycin to an OD_600_ of 0.6-0.8 at 37 °C, and then induced by addition of 660 μM isopropyl β-D-1-thiogalactopyranoside (IPTG) followed by growth for 16 h at 18°C. Cells were lysed using a microfluidizer and centrifuged for 60 min at 30,000 × g at 4 °C. The resulting soluble fraction was then applied to a Ni-Sepharose column (GE Healthcare Life Sciences) preequilibrated in buffer B (50 mM 3-(*N*-morpholino) propanesulfonic acid (MOPS) pH 7.5, 10% glycerol, 150 mM NaCl, 1 mM MgCl_2_, 1 mM 1 mM Tris(2-carboxyethyl)phosphine (TCEP)) with 15 mM imidazole. Buffer B containing 1 M NaCl was used to elute nucleic acid contaminants. Bound proteins were eluted using an imidazole gradient (50-500 mM) in buffer B over 10 CV. Fractions containing MRPP3 were pooled and TEV protease was added and the sample was dialyzed overnight against Buffer B at 4 °C. The sample was then applied to a second Ni-Sepharose column. The flow-through was collected, concentrated, and dialyzed into 50 mM MOPS (pH 7.5), 1 mM TCEP, 150 mM NaCl, and 10 % glycerol.

MRPP1 and MRPP2 are prepared using a pCDFDuet-1 co-expression vector (pHisTEVMP1+MP2) containing an N-terminal truncation of the MRPP1 gene (*TRMT10C*) corresponding to residues 40-403 (Δ39MRPP1) with an N-terminal His_6_-tag followed by a TEV protease site and the full-length MRPP2 gene (*SDR5C1*) (44). Rosetta (DE3) or Rosetta2 (DE3) *E. coli* cells were transformed with the pHisTevMP1+MP2 plasmid and grown to an OD_600_ of 0.6-0.8 in terrific broth media containing 50 mg/L streptomycin and 34 mg/L chloramphenicol at 37 °C. Expression was induced by addition of 0.5 mM IPTG and cells were grown at 18 °C for 16 h after induction. Cells were harvested and lysed using a microfluidizer in buffer A (20 mM Tris-HCl pH 7.5 or 50 mM MOPS pH 7.5, 1 mM TCEP, 10% glycerol, 1 M NaCl, 15 mM imidazole) and one tablet of Complete EDTA free protease inhibitor cocktail (Roche). Cell lysate was pelleted for 60 min at 30,600 × g at 4 °C. The soluble fraction was added to a 5 mL Ni-Sepharose fast-flow resin (GE Healthcare Life Sciences) pre-equilibrated with buffer A at room temperature. The column was washed with buffer A containing 150 mM NaCl and bound proteins were eluted by an imidazole gradient (50 mM to 500 mM). Fractions containing His_6_-MRPP1/2 were pooled and dialyzed overnight at 4 °C against buffer A containing 150 mM NaCl and His_6_-tagged TEV protease. The sample was applied to a second Ni-Sepharose fast-flow resin pre-equilibrated with buffer A containing 150 mM NaCl. The flow-through fractions containing MRPP1/2 were pooled and dialyzed overnight into dialysis buffer (50 mM MOPS pH 7.5, 300 mM NaCl, and 1 mM TCEP), and then concentrated using a centrifugal filter with a 10K MWCO (Amicon). The sample was loaded onto a Sephacryl-200 gel-filtration column (GE Healthcare) that was pre-equilibrated with the dialysis buffer. Protein was eluted at 1 mL/min and the A_280_ value was monitored. Peak fractions were analyzed by SDS-PAGE and coomassie blue stain.

### Pre-tRNA preparation

*B. subtilis* pre-tRNA^Asp^ was transcribed from a PCR template as previously described (2, 3) and fluorescently labeled at the 5’-end according to literature procedures (45). The concentration of pre-tRNA substrate was calculated by absorbance using an extinction coefficient of 685,000 M^−1^cm^−1^ at 260 nm. Labeling efficiency was assessed by fluorescein absorbance at 492 nm using an extinction coefficient of 78,000 M^−1^cm^−1^.

### Fluorescence polarization (FP) assay

The activity of PRORP1 catalyzed pre-tRNA cleavage was measured by monitoring the fluorescence anisotropy (FA) signal (λ_ex_ = 485 nm and λ_em_ = 535 nm) over time at 25 °C in buffer conditions optimized for a high-throughput format, 30 mM MOPS pH 7.8, 5 mM MgCl_2_, 100 mM NaCl, 1 mM TCEP, 500 nM (12 μg/mL) yeast tRNA, 5 mM spermidine, 6.65 μg/mL BSA, and 0.01% NP-40. Yeast tRNA^mix^ (Roche) was used to reduce the number of compounds that inhibit PRORP1 activity by targeting tRNA. Prior to each reaction, pre-tRNA was heated for 3 min at 95 °C to unfold the RNA and refolded in the reaction buffer at room temperature (25 °C) for 30 min. Reactions were performed in black, 96-well plates (Corning Incorporation, #3686) using a TECAN plate-reader (Infinite F500, G-factor = 1.05) at 25 °C and the FA values recorded. Reactions were initiated with the addition of substrate to enzyme. Experiments were done in the presence of excess substrate. The concentration of enzyme was 0.3-5 nM, the concentration of labeled substrate (Fl-pre-tRNA) was 40 nM, and the concentration of pre-tRNA was 120-4000 nM. The concentration of Fl-pre-tRNA was held constant in all reactions.

### FP data analysis

To correct for small fluctuations in the FA reading that are inherent in the instrument, a correction factor (CF) was calculated using a negative control well without the addition of enzyme.

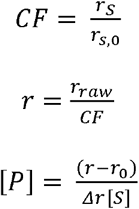

[P] = concentration of product

[S] = concentration of substrate

r = corrected FA value in millianisotropy (mA)

r_0_ = corrected FA at initial time

r_s_ = FA reading of substrate only control

r_s,0_ = FA reading of substrate only at initial time

r_raw_ = raw FA reading

Δr = total change in FA

### High-throughput screening

A small molecule library of compounds from microsources and pharmacologically active compounds (Prestwick, LOPAC, MS2400, BioFocus NCC, ~6,000 compounds) was screened at the Center for Chemical Genomics (CCG) at the University of Michigan. The FA cleavage assay was performed using 0.3 nM PRORP1, 800 nM pre-tRNA containing 40 nM Fl-pre-tRNA, at 22 °C in 30 mM MOPS pH 7.8, 5 mM MgCl_2_, 100 mM NaCl, 1 mM TCEP, 500 nM (12 μg/mL) yeast tRNA^mix^, 5 mM spermidine, 6.65 μg/mL BSA, 0.01% NP-40, 12.5 μM compound, and 1% DMSO. Enzyme is incubated with 25 μM compound or 1.25% DMSO in 384-well plates (Perkin Elmer Proxiplate 384F Plus #6008269) for 30 min before the addition of an equal volume of substrate (800 nM final concentration) to initiate the reaction. Each reaction was quenched with 80 mM CaCl_2_ after 1 h. The FA value was read on a PHERAstar plate reader and the Z-factor for each plate was calculated. The FA signal was used to detect any compounds that bound to pre-tRNA without PRORP1 present.

### Gel-based activity assay

To validate inhibitors identified from the high throughput FP assay (Fig. S4), we used an orthogonal gel-based assay to measure RNase P activity. This assay detects the presence of substrate and product by the fluorescence signal from fluorescein-labeled pre-tRNA run on a 20% PAGE gel containing 8 M urea. Compounds that resulted in no observed RNase P activity in the presence of 12.5 μM compound after 1 h were analyzed further using a dose response assay.

### Human mitochondrial RNase P (MRPP1/2/3) activity assay

The inhibition of human MRPP1/2/3 by gambogic acid and juglone was evaluated via dose-response using a quenched assay. Human MRPP1/2/3 activity was measured as a function of gambogic acid and juglone concentration using 50 nM MRPP3, 500 nM MRPP1/2, 500 nM pre-tRNA containing 40 nM Fl-pre-tRNA in 50 mM MOPS (pH 7.8), 4.5 mM MgCl_2_, 100 mM NaCl, and 1 mM TCEP (pH 7.5). Proteins were mixed together on ice and incubated at room temperature for 5 min before addition to pre-folded substrate at 37 °C to initiate the reaction. Substrate was folded as described in the FA assay with PRORP1. Reaction time points were taken and quenched with equal volumes of an EDTA-dye quench solution containing 100 mM EDTA, 0.05% xylene cyanol dye, 0.05% bromophenol blue dye, and 8 M urea. Samples were separated by 20% Urea PAGE (Ultra Pure SequaGel), fluorescence signal was quantified using ImageJ, and the data was analyzed using Prism. The SEM for the data is reported from the fit.

### *B. subtilis* RNase P activity assay

The inhibition of *B. subtilis* RNase P activity by gambogic acid and juglone was measured using a previously developed FP assay in real-time.^25^ Reactions were performed under MTO conditions using 20 nM RNase P and 500 nM Fl-pre-tRNA^Asp^ in 50 mM Tris (pH 7.2), 5 mM MgCl_2_, 100 mM KCl, 500 nM (12 μg/mL) yeast tRNA^mix^, 20 mM dithiothreitol (DTT), 10 mM spermidine, and 0.01% NP-40 at 37 °C.

### Dose-response and determination of mode of inhibition

The dose response was measured using the FP assay as described above with varying concentrations of compound. Compounds were incubated with PRORP1 at twice the reaction concentration for 30 minutes prior to the addition of equal volumes of substrate to initiate the reaction. Experiments were done under multiple turnover conditions (same buffer conditions as the FP assay). Each reaction was performed once to determine the initial velocity and the results were converted to activity (%) using DMSO as a 100% activity control and CaCl_2_ or no enzyme as 0% activity controls. PRORP1 activity was determined as a function of inhibitor concentration, and equation 1 was fit to the data to determine the *IC*_50_ value. The standard error of the mean (SEM) is reported from the fit.

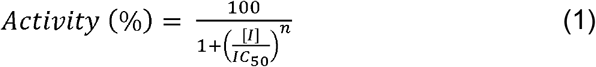

To identify the mode of inhibition of gambogic acid, PRORP1 activity was measured with varying pre-tRNA and gambogic acid concentrations. The Michaelis-Menten equation (Equation 2) was fit to each curve individually to calculate the apparent turnover rate constant (*k*_cat,app_) and apparent substrate concentration at half maximal activity (*K*_M,app_). Equations 2, 3, and 4 were also used in a global fit analysis on these data.

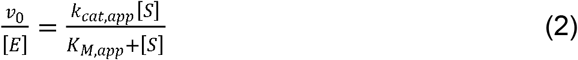

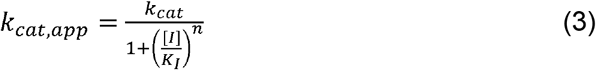

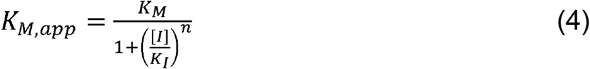

For the analysis of juglone, PRORP1 activity was measured with varying pre-incubation time and juglone concentration. Varying concentrations of juglone were pre-incubated with a 35 μL of PRORP1 from 2 min to 4 h in 30 mM MOPS pH 7.8, 5 mM MgCl_2_, 100 mM NaCl, 1 mM TCEP, 500 nM (12 μg/mL) yeast tRNA^mix^, 5 mM spermidine, 6.65 μg/mL BSA, 0.01% NP-40. The concentration of DMSO in each reaction was held constant at 1%. Reactions were initiated by addition of 5 μL of substrate for a final working concentration of 0.3 nM PRORP1 and 250 nM pre-tRNA, a substrate concentration well below the *K*_M_ value. The dependence of the initial velocity on the pre-incubation time was fit to a single exponential decay equation (Equation 5) to determine *k*_inact,app_ at each juglone concentration. The values for *k*_inact,app_ for various juglone concentrations (represented by [I]) were fit with Equation 6 to determine *k*_inact_/*K*_I_, assuming a 2-step inactivation mechanism.

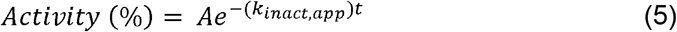

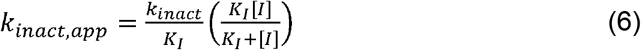

### Crystallization, Data processing and Refinement

Δ76-PRORP1 was expressed and purified for crystallization as described above. Crystals were grown in 0.1 M sodium citrate pH 5.5 and 18 % (wt/vol) PEG 3350 at 4 °C by sitting drop method with a 2:1 ratio of protein and reservoir solution.^19^ Δ76-PRORP1 crystals were soaked in reservoir solution containing (18% PEG 3,350, 0.1 M sodium citrate, pH 5.5, and 10 μM juglone), a saturating concentration of juglone (>10 μM) and incubated for 45 min, 3 h, and overnight in the presence or absence of 50 mM MnCl_2_ before harvesting. Harvested crystals were cryoprotected by soaking in 18% PEG 3,350, 0.1 M sodium citrate tribasic (pH 5.5), 20% (v/v) glycerol and 10 μM juglone for a few minutes prior to flash cooling in liquid N_2_. Diffraction data were collected on beamline GM/CA-CAT 23 ID-D and 23 ID-B with a Dectris Pilatus3 6M and a Dectris Eiger X 16M detectors respectively at Advanced Photon Source, Argonne National Laboratory (Argonne, IL). The data were indexed and processed with XDS and then reduced with aimless (46). Phaser MR was subsequently used for molecular replacement (47). Original models were subject to restrained refinement with phenix.refine (48). Refinement included simulated annealing in torsional and Cartesian space, coordination minimization, and restrained individual B-factor adjustment with maximum-likelihood targets. Energy-minimized coordinates and a cif formatted topology library for juglone were obtained by ProDrg webserver (49). Refmac5 in the CCP4i suite was subsequently employed for restrained refinement model building and modification with Coot (50, 51). Crystallographic information and refinement statistics are provided in Table S3. The geometric quality of the model and its agreement with the structure factors were assessed with MolProbity (52). Figures showing crystal structures were generated in PyMOL (53). Atomic coordinates and structure factors for PRORP1 with juglone have been deposited in the Protein Data Bank (PDB).

## Data availability

Structural data are deposited in the RCSB Protein Data Bank. PDB-ID-codes: 6BV5 (DOI: https://doi.org/10.2210/pdb6BV5/pdb), 6BV6 (DOI: https://doi.org/10.2210/pdb6BV6/pdb), 6BV8 (DOI: https://doi.org/10.2210/pdb6BV8/pdb), and 6BV9 (DOI: https://doi.org/10.2210/pdb6BV9/pdb).

## Supporting Information

This article contains supporting information

## Acknowledgements

We thank Dr. Xin Liu for her expertise in developing RNase P high throughput screening assays and Martha Larsen for her assistance in our high throughput screening work at the Center of Chemical Genomics at the University of Michigan. We also would like to thank Dr. Albert Epshteyn, Dr. Bradley Klemm and Dr. Danielle Zurcher for advice and discussions on assessing a variety of juglone-cysteine conjugates in the PRORP1 structures. We would like to thank the National Institute of General Medical Sciences and National Cancer Institute Collaborative Access Team (GM/CA CAT) at the Advanced Light Source for beam time. This work was supported by the National Institutes of Health [GM55387 to C.A.F.] and [GM117141 to M.K.] and the Robert A. Welch Foundation [A-1987 to C.A.F.].

## Conflict of Interest

The authors declare that they have no conflicts of interest with the contents of this article.

## Notes

### Competing Interest Statement

The authors have declared no competing interest.

