## Supplemental Information for "Mode of inhibition of RNase P by gambogic acid and juglone"

**Supplementary data**

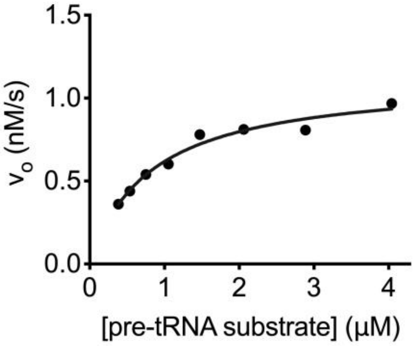


**Supplementary Figure 1.** PRORP1 activity was measured using the FP assay using 5 nM PRORP1 and varying pre-tRNA substrate (0.38 – 4 µM) in 30 mM MOPS pH 7.8, 5 mM $MgCl_2$, 100 mM NaCl, 1 mM TCEP, 500 nM (12 µg/mL) yeast $tRNA^{mix}$, 5 mM spermidine, 6.65 µg/mL BSA, and 0.01% NP-40 at 22 °C. The Michaelis-Menten equation was fit to the data to determine $k_{cat}$ (0.22 ± 0.01 $s^{-1}$) and $K_M$ (0.8 ± 0.1 µM) with error given as standard error of fit.

**Table 1. List of 42 compound hits that decrease PRORP1 activity more than 2.5-fold at 12.5 µM[1]**

| | |
|---|---|
| 1-aminoindan-1,5-dicarboxylic acid | erythrosine sodium |
| 1-benzyloxycarbonylaminophenethyl chloromethyl ketone | evans blue |
| S-2-(3-aminopropylamino)-ethylphosphorothioic acid | gambogic acid |
| 2-[3-(1,3-Dihydro-1,3,3-trimethyl-2H-indol-2-ylidene)-1-propenyl]-3-ethyl-benzothiazolium iodide | garcinolic acid |
| 2E-N-[(5-bromo-2-methoxyphenyl)sulfonyl]-3-[2-(2-naphthalenylmethyl)phenyl]-2-propenamid | gentian violet |
| 3-[3,5-dibromo-4-hydroxylbenzoyl]-2-ethylbenzofuran | hexachlorophene |
| 5-(1-aziridinyl)-2,4-dinitrobenzamide | juglone |

[1] PRORP1 activity was measured using the high throughput FP assay using 0.3 nM PRORP1, 800 nM pre-tRNA containing 40 nM Fl-pre-tRNA, at 22 °C in 30 mM MOPS pH 7.8, 5 mM $MgCl_2$, 100 mM NaCl, 1 mM TCEP, 500 nM (12 µg/mL) yeast $tRNA^{mix}$, 5 mM spermidine, 6.65 µg/mL BSA, 0.01% NP-40, 12.5 µM compound, and 1% DMSO. Compounds were incubated with PRORP1 at twice the reaction concentration for 30 minutes prior to the addition of equal volumes of substrate to initiate the reaction. Each reaction was performed once to determine the initial velocity and the results were converted to activity (%) using DMSO as a 100% activity control and $CaCl_2$ or no enzyme as 0% activity controls.

| | |
|---|---|
| 5-iodo-3-[(3,5-dibromo-4-hydroxyphenyl)methylene]-2-indolinone | laphachol methyl ether |
| agaric acid | lobaric acid |
| anthothecol | methyl gambogate |
| antiarol | oleic acid |
| aurintricarboxylic acid | phenethyl caffeate |
| beta-lapachone | phenulmercuric acetate |
| betulinic acid | plumbagin |
| bexarotene | pyrithione zinc |
| bithionol | pyrvinium pamoate |
| cearoin | sanguinarium chloride |
| cedrelone | sodium tetradecyl sulfate |
| chloranil | stigmasta-4,22-dien-3-one |
| citropten | tetrahydrogambogic acid |
| dihydrogambogic acid | tetrachloroisophthalonitrile |

**Table 2. PRORP1 activity at 6.25 µM of inhibitor[2].**

| Compound name | *Activity (%)* |
|---|---|
| anthothecol | 15.5 |
| aurintricarboxylic acid | 46.5 |
| 1-benzyloxy carbonyl amino phenylethyl chloromethyl ketone | 19.9 |
| chloranil | 40.2 |
| evans blue | 20.2 |
| gambogic acid | 29.1 |
| gentian violet | 0 |
| juglone | 20.2 |
| phenylmercuric acetate | 32.2 |
| plumbagin | 2.6 |
| methyl gambogate | 34.8 |

[2] PRORP1 activity was measured using the FP assay at four to six inhibitor concentrations using 0.3 nM PRORP1, 800 nM pre-tRNA containing 40 nM Fl-pre-tRNA, at 22 °C in 30 mM MOPS pH 7.8, 5 mM $MgCl_2$, 100 mM NaCl, 1 mM TCEP, 500 nM (12 µg/mL) yeast $tRNA^{mix}$, 5 mM spermidine, 6.65 µg/mL BSA, 0.01% NP-40, 12.5 µM compound, and 1% DMSO. Compounds were incubated with PRORP1 at twice the reaction concentration for 30 minutes prior to the addition of equal volumes of substrate to initiate the reaction. Each reaction was performed once to determine the initial velocity and the results were converted to activity (%) using DMSO as a 100% activity control and $CaCl_2$ or no enzyme as 0% activity controls.

**Table 3. X-Ray Crystallography Data Collection and Refinement Statistics**

| | PRORP1/Mn + Juglone (45 min soaks) | PRORP1 + Juglone (3 hour soaks) | PRORP1/Mn + Juglone (3 hour soaks) | PRORP1 + Juglone (overnight) |
|---|---|---|---|---|
| **Data collection** | | | | |
| Beamline | APS, GMCA 23-IDD | APS, GMCA 23-IDD | APS, GMCA 23-IDD | APS, GMCA 23-IDB |
| Wavelength (Å) | 1.0332 | 1.0332 | 1.0332 | 1.0332 |
| Resolution (Å) | 50.0-1.79 (1.83-1.79) | 50.0-2.20 (2.26-2.20) | 50.0-2.10 (2.16-2.10) | 50.0-2.10 (2.16-2.10) |
| Space group | $P2_12_12_1$ | $P2_12_12_1$ | $P2_12_12_1$ | $P2_12_12_1$ |
| Cell dimensions (Å) | a = 41.6, b = 110.8, c = 140.7 | a = 41.7, b = 112.2, c = 139.2 | a = 41.7, b = 112.2, c = 139.2 | a = 41.8, b = 111.8, c = 139.3 |
| Cell dimensions (°) | α = β = γ = 90 | α = β = γ = 90 | α = β = γ = 90 | α = β = γ = 90 |
| $R_{sym}$ (%) | 3.6 (47.8) | 4.0 (69.2) | 4.8 (63.6) | 5.5 (71.7) |
| <I/σ> | 15.9 (1.9) | 11.2 (2.1) | 13.4 (1.6) | 17.5 (2.2) |
| CC(1/2) | 1.0 (0.75) | 1.0 (0.74) | 1.0 (0.56) | 1.0 (0.75) |
| Multiplicity | 3.7 (3.9) | 3.4 (3.3) | 3.5 (3.4) | 6.7 (6.5) |
| Completeness (%) | 99.1 (96.3) | 98.1 (95.8) | 99.2 (97.9) | 100 (99.7) |
| **Refinement** | | | | |
| Resolution range | 87.1-1.79 | 87.3-2.20 | 87.0-2.10 | 87.2-2.10 |
| Number of reflections (work/test set) | 57980/3033 | 31891/1637 | 36554/1900 | 37078/1982 |
| $R_{work}/R_{free}$ (%) | 19.6/22.3 | 20.6/23.6 | 20.0/24.7 | 19.6/24.2 |
| No. of atoms | | | | |
| Protein | 3769* | 3755* | 3758* | 3760* |
| Water | 239 | 118 | 173 | 193 |
| Modified Cysteines | 19 | 57 | 38 | 57 |
| Zn/Mn | 1/1 | 1/0 | 1/2 | 1/0 |
| B-factors ($Å^2$) | | | | |
| Protein | 48.4 | 59.0 | 59.2 | 62.4 |
| Water | 42.0 | 43.7 | 51.5 | 50.7 |
| Modified Cysteines | 53.2 | 77.5 | 71.0 | 78.4 |
| Zn/Mn | 39.7/32.0 | 45.8/na | 52.8/63.2 | 51.3/na |
| Rmsd deviations | | | | |
| Bond lengths (Å) | 0.010 | 0.010 | 0.011 | 0.012 |
| Bond angles (°) | 1.28 | 1.35 | 1.39 | 1.45 |
| Ramachandran plot | | | | |
| Favored/allowed/outliers | 98.3/1.7/0.0 | 97.8/2.0 /0.2 | 98.1/1.7/0.2 | 98.0/2.0/0.0 |
| MolProbity Score | 0.64 ($100^{th}$ percentile) | 0.93 ($100^{th}$ percentile) | 1.00 ($100^{th}$ percentile) | 0.97 ($100^{th}$ percentile) |
| PDB | 6BV5 | 6BV6 | 6BV8 | 6BV9 |

* The modified cysteines were not counted

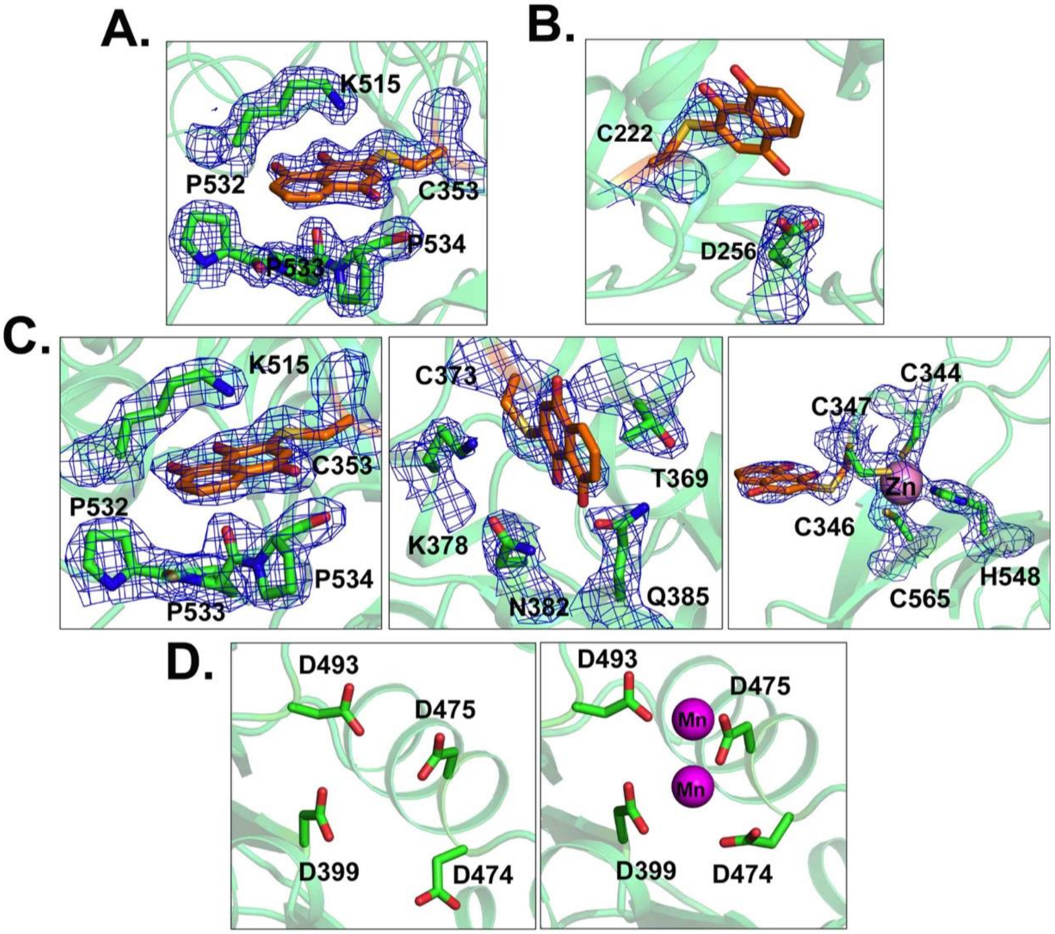


**Supplementary Figure 2**. (A) Electron densities of juglone conjugated cysteine (JCC) 353 and nearby amino acids in PRORP1 crystals soaked in juglone for 45 minutes (PDB ID: 6BV5). (B) Electron densities of JCC 222 and nearby amino acid in PRORP1 crystals soaked in juglone for overnight (PDB ID: 6BV9). (C) Electron densities of residues JCC 353, 373, 346 and nearby amino acids in PRORP1 crystals soaked in juglone for 3 hours (PDB ID: 6BV6). (D) Active site residues of PRORP1 in the absence (first panel) and presence (second panel) of active site metal ($Mn^{2+}$) in 3 hours juglone soaked crystals (PDB IDs: 6BV8).

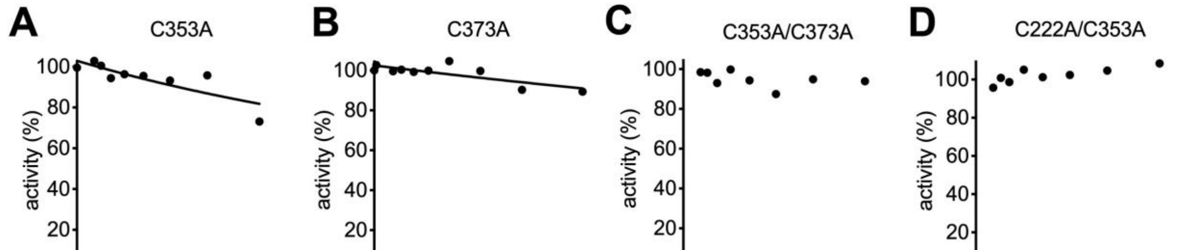


**Supplementary Figure 3.** PRORP1 activity was extrapolated to an incubation time of zero and plotted against juglone concentration to determine $K_I$. Single cysteine mutations increased the $K_I$ values (>10 μM) while inhibition at time 0 was not observed with the double-cysteine mutations. Reactions were performed with 0.3 nM PRORP1 variant, 250 nM pre-tRNA containing 40 nM Fl-pre-tRNA, at 22 °C, in 30 mM MOPS, pH 7.8, 1 mM TCEP, 5 mM $MgCl_2$, 500 nM (12 μg/mL) yeast tRNA, 5 mM spermidine, 6.65 μg/mL BSA, and 0.01% NP-40.

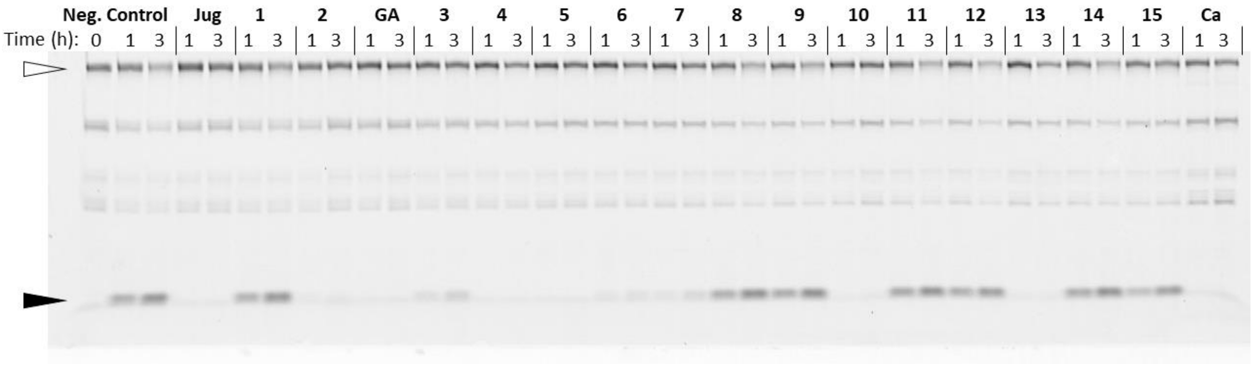


**Supplementary Figure 4.** A PRORP gel inhibition assay with 0.30 nM PRORP and 800 nM fluorescently labeled *B. Subtilis* pre-$tRNA^{Asp}$. Assays were carried out under the same conditions as the fluorescent HTS described in Materials and Methods. PRORP was incubated with 12.5 μM of each compound prior to reaction initiation by the addition of pre-tRNA. The reaction was quenched at 1 and 3 hour time points with an equal volume of 100 mM EDTA, 6 M urea, 0.1% bromophenol blue, 0.1% xylene cyanol. The 5'-leader product (closed arrow) was resolved from pre-tRNA substrate (open arrow) on a denaturing 20% polyacrylamide gel. The compound in each lane is indicated as follows: (Neg. Control) no inhibitor control, (Jug) juglone, (1) cearoin, (2) anthothecol, (GA) gambogic acid, (3) tetrahydrogambogic acid, (4) agaric acid, (5) 1-benzyloxycarbonylaminophenethyl chloromethyl ketone, (6) chloranil, (7) aurintricarboxylic acid, (8) phenethyl caffeate, (9) pyrvinium pamoate, (10) L-798106, (11) CB 1954, (12) AIDA, (13) GW5074, (14) AC-93252 iodide, (15) bexarotene and (Ca) 20 mM $Ca^{2+}$ in absence of $Mg^{2+}$ as a positive inhibition control.

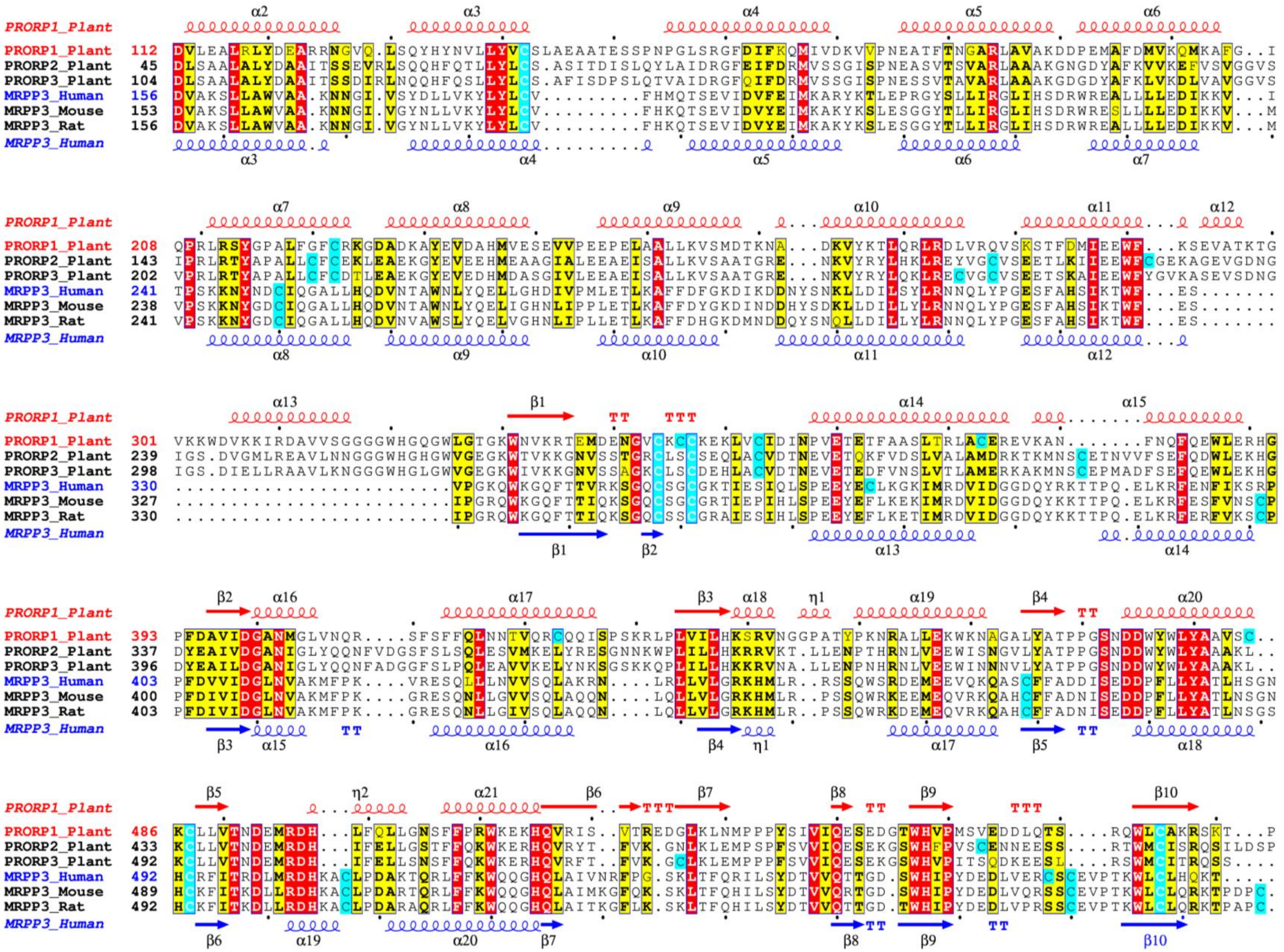
PRORP1_Plant
PRORP1_Plant
PRORP2_Plant
PRORP3_Plant
MRPP3_Human
MRPP3_Mouse
MRPP3_Rat
MRPP3_Human

**Supplementary Figure 5.** A multiple sequence alignment of PRORP proteins. Aligned are three *Arabidopsis thaliana* PRORP paralogs and three mammalian MRPP3 homologs from *Homo sapiens*, *Mus musculus* and *Rattus norvegicus*. Secondary structure for *A. thaliana* PRORP1 (PDB) and *H. sapiens* MRPP3 (PDB) is annotated above and below the amino acid sequence, respectively. Conserved amino acids are highlighted in red, chemically similar residues bold and boxed in yellow, cysteine residues highlighted in cyan.
